# The effect of auditory stimulation on the nonlinear dynamics of heart rate: the impact of emotional valence and arousal

**DOI:** 10.1101/2022.03.19.484969

**Authors:** Dmitri Dimitriev, Olga Indeykina, Aleksey Dimitriev

## Abstract

**Background:** Although it is known that sound exposure evokes changes in autonomic activity, the effects of noise and music on the nonlinear behavior of heart rate fluctuations remain poorly understood and controversial. This study aims to assess the influence of sound subjective emotional valence and arousal on the nonlinear characteristics of the autonomic nervous system during passive listening.

**Methods:** In this study, forty-two subjects listened to four sounds: (1) white noise, (2) road traffic noise, (3) excitatory music, and (4) a lullaby. The experiment consisted of two consecutive sessions: five minutes of rest, followed by five minutes of listening. RR intervals were recorded during both sessions. The following linear and nonlinear heart rate variability indices were computed: SDNN, RMSSD, F, HF, approximate (ApEn) and sample entropy (SampEn), correlation dimension (D2), Poincare plot indices (SD1, SD2), fractal scaling exponents (alpha1, alpha2), and recurrence plot indices (Lmean, Lmax, DET, LAM, Vmax, TT, ShanEn).

**Results:** Excitatory music was associated with significant decrease in SDNN and RMSSD, diminished HF, and a substantial reduction in LF. Excitatory music exposure induced significant increases in DET, SD1 and SD2, but changes in DFA, SampEn, and D2 were nonsignificant. Traffic noise, white noise, and the lullaby did not cause significant changes in the measures of heart rate variability.

**Conclusion:** Presentation of music evoking strong negative emotions elicits a prominent decrease in respiratory sinus arrhythmia. Poincare plot and recurrence plot measures possess high sensitivity to high arousal and unpleasant music. Contrary to previous studies, we did not find the effects of relaxing music on heart rate variability.

## Introduction

When considering music in terms of psychology and physiology, it delves into the relationship between mind and body. Merriam [1] identified ten music functions, the first of which was expressing and evoking emotions.

Although music philosophers debate the nature of music-evoked emotions, there is evidence that music can directly induce changes in the significant components of emotion, including subjective feeling and physiological arousal (autonomic and endocrine changes) [2]. Due to functional interconnections between brain structure involved in the auditory processing and limbic system, music can evoke changes in the activity of brain regions underlying emotions. Moreover, the auditory system contributes to various neural circuits that govern autonomic and somatic motor behavior expression. In accordance with general theories of emotion, arousal is a crucial component of emotional responses to music [3]. Currently, arousal is defined as the degree of excitement or motivational activation [4; 5] a person experiences as a reaction to emotional stimuli. Valstar [6] denotes arousal as a global feeling of dynamism or lethargy that involves mental activity and physical preparedness to act.

The association between emotional arousal and activity of the sympathetic branch of the autonomic nervous system has been well-established [7]. Anger and anxiety stimuli evoke an increase in heart rate, respiratory rate, and blood pressure. Based on this effect, one could expect that music with low anxiety induces a shift of autonomic balance towards dominance of the parasympathetic nervous system, but the reality is more complex and contradictory. Pérez-Lloret et al. [8] assessed emotional and autonomic reactions to different “relaxing” music styles and found that preferred “new age” music-induced significant reduction of HF spectral component and increase of LF/HF ratio but the absence of changes in heart rate. With some reservations, these results point to a shift in autonomic balance in a setting with a low parasympathetic nervous tone [9]. Other investigators observed, however, that relaxed music significantly increases vagal activity and can even be used to heal.

Intense emotional responses to music could be understood in terms of very high levels of physiological arousal. Iwanaga and Tsukamoto reported that excitative music evokes significant depression of amplitude of high-frequency variance of heart rate [10]. In research of self-reported emotional responses to the music, it was found that state arousal levels were higher in subjects listening to heavy metal than in subjects exposed to other genres, including pop music, classical music, and country [11]. According to the current theory, one would expect prominent activation of the sympathetic branch during heavy metal listening [12], but investigations of the causal relationship between heavy metal music exposure and ANS activity have yielded contradictory findings. In a study by Nater et al. [13], skin conductance reactivity and heart rate were significantly higher during heavy metal than during listening to relaxing music. Kalinowska et al. [13] found an absence of significant differences between heart rate and blood pressure before and after listening to heavy metal music.

Another essential feature of emotion is valence, and all music stimuli may be classified along this dimension as a positive or negative experience [15]. Nyklíček et al. [16] demonstrated that musically induced emotions (happiness, sadness, serenity, and agitation) might be discriminated in terms of autonomic activity measures.

Traffic noise is an essential part of the urban sound landscape. Recio et al. [17] presented an integrative stress model linking environmental noise exposure with noise with cardiovascular, respiratory, and metabolic disorders and diseases. According to this model, autonomic response to noise exposure is the critical element of noise-induced stress. Heart rate variability (HRV) indexes the activity of integrated regulatory systems, which operate on different time scales; HRV reflects an adaptation of the autonomic nervous system to different environmental and internal challenges. Daytime traffic noise exposure induced an immediate increase in HR and changes in HRV indices (decrease in LF and HF), associated with equivalent continuous sound pressure levels [18]. The record of noise level and heart rate variability with a personal dosimeter and an electrocardiography sensor on the chest reveals a significant negative correlation between an increase in sound pressure level and heart rate variability measures [19]. White noise listening evoked a significant increase of LF and LF/HF in healthy young adults [20], and a comparison of different sound level reveal that HF for sine wave sound was significantly greater than that for white noise [21]. Natural sounds are coupled with affective emotions and unnatural sounds, like pure tone and white noise, used to eliminate or minimize the association of sounds with real-life events [19].

The loudness of sound stimuli is associated with aversive emotions that sounds with a high-pressure level (around 100 dB) are often used to induce strong negative emotions [23]. In daily life, one is unlikely to encounter 100 dB sounds routine; sounds at approximately 50 dB are most commonly heard, and most human conversations and mechanical noise occur at moderate loudness levels [24].

Acoustic stimuli with high-pressure levels induce concurrent activation vestibular processes that directly affect autonomic balance and changes in higher auditory cognitive processes [25]. At moderate sound level, vestibular activation of SNS does not play a significant role in autonomic reaction to sound exposure [26].

Most research has focused on linear measures of autonomic activity, such as time and frequency domain markers of heart rate variability. However, the experimental results and mathematical models support the view that autonomic rhythm reflects the forcing of a nonlinear oscillator rather than periodic inhibition of unstructured, random activity [27]. Physiological mechanisms underlying heart rate variability include stochastic processes at the cellular level, the influence of respiration on the heart rate, and interactions of the multiple feedback loops regulating the cardiovascular system. Hence, neurocardiac regulation may be nonlinear in structure due to the multi-level nature of the interactions between the ANS and the various control mechanisms [28]. The term nonlinear describes a comprehensive class of systems (including physiological), where the output is not linearly dependent on the strength of the applied stimulus. Evidence for nonlinearities in cardiac physiology includes abrupt changes (bifurcations), self-sustained and complex oscillations, chaotic behavior, fractal structures, and hysteresis. Numerous algorithms have been developed to describe the dynamic fluctuations in heart rate data, and a variety of nonlinear methods have been used to analyze RR interval dynamics.

Heart rate complexity arises from the components of intrinsic system dynamics, especially from the nonlinear interplay of different physiological feedback loops [28]. The most widely used set of HRV complexity measures is based on the concept of entropy. Several entropy-based complexity assessment methods have been proposed, although there is no direct correspondence between entropy and mathematical complexity [29]. Pincus [30] created the approximal entropy (ApEn) measure. To overcome its limitations, ApEn was later improved and termed “sample entropy” (SampEn) by Richman and Moorman [31]. Entropy-based measures appoint the highest values to uncorrelated random signals, which are very unpredictable but not “complex.”

Chaos is the essential characteristic of nonlinear systems, including the ANS [32]. In the presence of chaos, the complexity of the heart rate attractor can be quantified via correlation dimension (D2). D2 measures signal complexity and reflects the number of underlying functional mechanisms responsible for a beat-to-beat time series [33].

The recurrence of states is a fundamental characteristic of dynamic systems and is typical for systems with chaotic dynamics. The method of recurrence plots (RP) was developed to visualize the dynamics of phase space trajectories [34]. For quantifying the structures yielded by RPs, several measures of complexity (based on recurrence point density and the diagonal and vertical line structures) have been proposed [34]. These measures are known as recurrence quantification analysis (RQA). Several studies have used RQA in physiological signal analysis, which has deepened the understanding of healthy dynamics and pathological disturbances.

Quantitative analysis of heart rate fluctuations reveals the scale-invariant (fractal) behavior of RR intervals. Detrended fluctuation analysis (DFA) describes fractal scaling properties and the long-range correlations in noisy, nonstationary time series, avoiding the spurious results that are possibly caused by non-stationarity. DFA provides a short- and long-term exponent (termed α1 and α2) to characterize the degree of correlation among time scales [35].

HRV is the product of complex neuroarchitecture, with different links between cortical, midbrain, and brainstem structures [36]. Listening to music evokes complex EEG activity with nonlinear dynamic properties [37]. The affective sounds and pictures elicit changes in the nonlinear dynamics of RR intervals [38]. Theoretically, we can assume that music should be able to modify HRV nonlinear dynamics.

This study aimed to determine potential changes in nonlinear HRV indices during passive listening to sounds. In addition, the present study sought to evaluate whether emotional valence and arousal contributed to the changes in linear and nonlinear measures of HRV.

## Materials and Methods

An a priori sample-size calculation with the software G*Power [39] indicated that 35 participants would be sufficient to detect a medium-size effect (0.5) with a power of 0.8. The final sample included 42 female participants between 19 and 24 years (mean age = 21.2 ± 1.). All subjects were healthy without neurological, psychiatric, cardiovascular, autonomic, or audiological disorders. All subjects denied consuming any medication. The study protocol confirmed to the Declaration of Helsinki principles, it was approved by the local ethics committee. Volunteers provided written, informed consent before participation.

Four sound stimuli were used: (1) road traffic noise (recorded at a busy street crossing), (2) white noise (generated using SoundForge 7.0), (3) excitatory music (Diamanda Galas’ Women with Steak Knives), and (4) a lullaby Ba Mo Leanabh (O My Baby) (Arr. William Jackson and Mackenzie, Fiona McKenzie).

These sounds were chosen according to a procedure described by Lin et al. [40]. In a pilot study, 70 students estimated ten musical pieces according to valence and arousal. Based on the evaluations, the above-mentioned sounds were chosen because of the variety in terms of arousal and valence ratings. The sounds were presented using the Yamaha HiFi system (PianoCraft E810). Playback settings were established beforehand so that the sound pressure level was 70 dBA (sound level meter Center 320, Taiwan). The stimulus duration was set to five minutes.

The affect grid [41] was used for the assessment of emotional reactions to the sounds. The affect grid comprises a number of squares that divided the two-dimensional space, with the horizontal axis identified as valence and the vertical axis as arousal. Each square depicted a combination of valence (−4 = ‘strong negative valence’ and 4 = ‘strong positive valence’) and emotional arousal (−4 = ‘very low arousal’ and 4 = ‘very high arousal’). The research design divided the experiment into four blocks of even length, with each experimental block consisting of rest and listening stages. Each stage lasted five minutes, using one-minute intervals. During the entire experimental session, the participants sat in a comfortable armchair. An electrocardiogram (ECG) signal was sampled at 1 kHz during both experiment stages. Successive RR intervals of the participants were obtained from the ECG recordings to calculate the HRV. The sequences of RR intervals were analyzed using the RHRV analysis package [42].

From each HRV series, several time domain and frequency domain indices were calculated:

the standard deviation of NN intervals (SDNN),
the square root of the mean of the sum of the squares of the differences between subsequent NN intervals (RMSSD),
the power calculated within the LF and HF bands.

The nonlinear behavior of heart rate could be represented by a trajectory through multidimensional space (phase space). The subset of the phase space corresponding to the typical behavior of the dynamical system is known as the attractor. In this article, the correlation dimension (D2) and entropy measure (SampEn) were used to describe the complexity of the attractor.

Fractal scaling exponents of heart rate variability were analyzed using the DFA method. The scaling exponent, α, represented the slope of the line related to the double log plot of the fluctuation function, F(n), and the size of the window, n. The short-term fluctuation slope (α1) was calculated for a period of four to 11 beats, and long-term fluctuations (α2) were calculated for periods more extended than 11 beats [35].

The Poincaré plots, correlating RRn on the x-axis and RRn +1 on the y-axis, were used to investigate HRV as a series of discrete events and beat-to-beat cycles. Three indices were calculated from the Poincaré plots: standard deviation of the short-term RR-interval variability (width of the cloud, SD1), the standard deviation of the long-term RR-interval variability (length of the cloud, SD2), and the ratio of SD2/SD1.

Recurrence plots are a graphical method for visualizing dynamical systems’ recurrence states and estimating time-series complexity. The primary step in recurrence analysis is the computation of the N × N matrix elements according to equation 1 as follows:

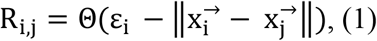

where N is the number of states, ε_i_ is a threshold distance, ∥·∥ is a norm, and Θ is the Heaviside function. The horizontal coordinate, i, in the RP relates to the system’s state at i, and each vertical coordinate, j, relates to the state at j. Points were denoted as recurrent (black point) whenever the distance between paired vectors, 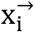 and 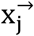, was below a threshold distance. Stochastic behaviour causes no (or short) diagonals, whereas deterministic behaviour causes longer diagonals and less single, isolated recurrence points.

The embedding dimensions, m, and time delay, τ, were estimated for each data set using the mutual information function and the false nearest neighbor method, respectively. The phase space was reconstructed using the values m = 10 and τ = 1, in accordance with previous studies. In the present study, the Euclidean method was employed for calculating the distances between individual points. The following measures were derived from RP: mean line length (Lmean), maximum line length (Lmax), recurrence rate (REC), determinism (DET), laminarity (LAM), trapping time (TT), maximal vertical length (Vmax), and Shannon entropy of line length distribution (ShanEn). The REC is the most straightforward measure of the RP and is shown as the black dots in the RP. A higher level of recurrence rate in RP (higher %Rec) can itself produce more extended lengths of diagonal lines. The threshold distance, ε, was set to achieve 5% of recurrent points for each participant’s resting and listening phase [43]. Lmean is the average diagonal line length. Determinism is the fraction of recurrence points forming diagonal lines of minimal length, and this measure indicates the predictability of the system. It has been showed that the maximum line length, denoted Lmax, and it’s inverse (the divergence (DIV)) are associated with the largest Lyapunov exponent (LLE) [44]. LAM of a recurrence plot is a fraction of recurrence points that form vertical lines. It reveals chaos-chaos transitions in short and stationary time series. Maximum vertical length (Vmax) reveals information about the time duration of the laminar states. Trapping time (TT) is the mean length of vertical lines in the recurrence plot. The TT represents the length of time that the dynamics remain trapped in a specific state. Shannon entropy of the distribution of the diagonal line lengths measures the complexity of the deterministic structure in the system [45].

Freidman ANOVAs were used to determine whether the effects of sound exposure differed from one another in terms of valence and arousal. Wilcoxon matched-pairs signed-ranks tests were used to determine the significance of differences between HRV measurements. Data are mean ± SE.

## Results

Figure 1 shows that subjects were exposed to an equivalent sound level approximately between 70 dB and 80 dB.

**Fig. 1.**
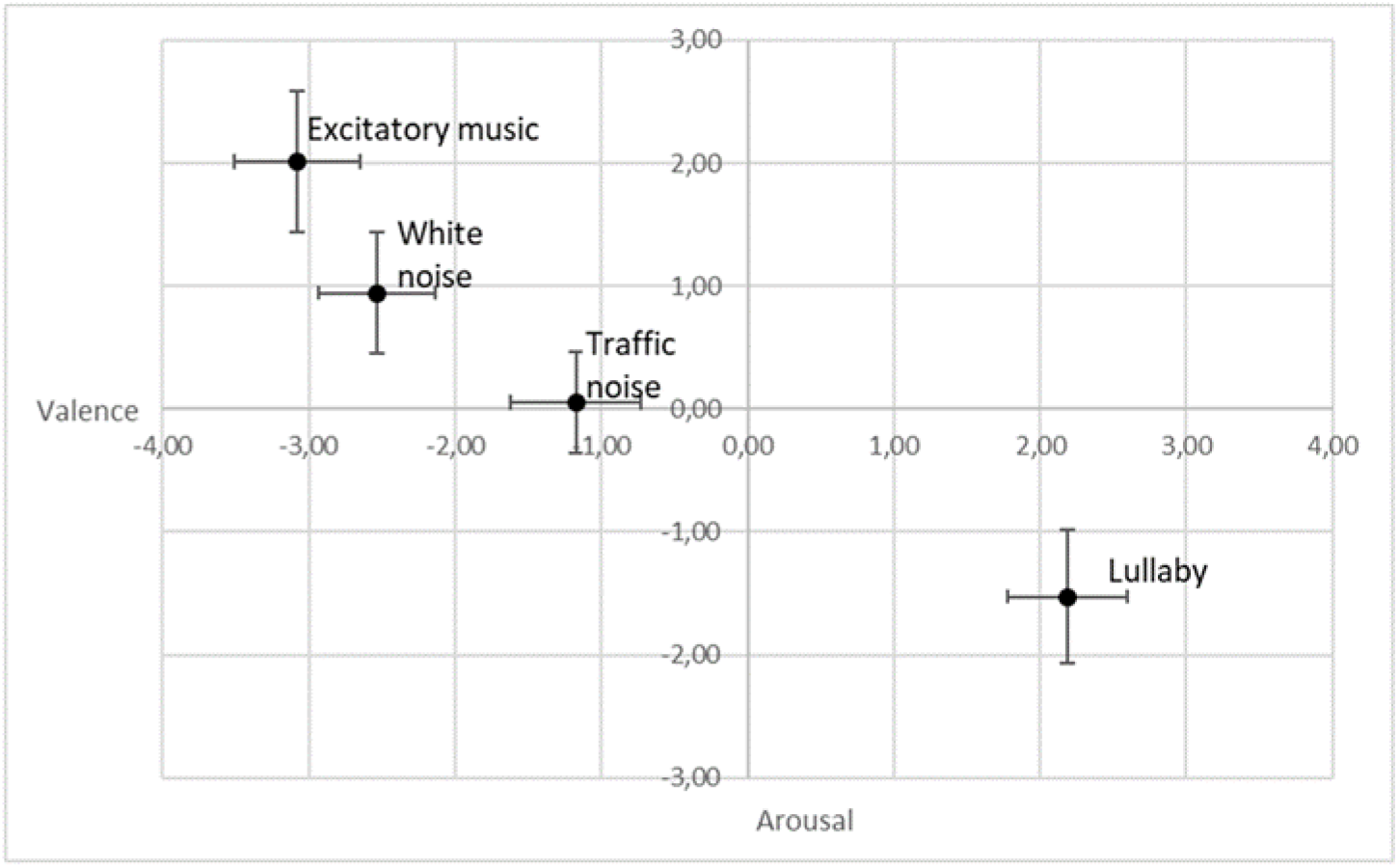
Sound ratings in the affect grid. Filled circles indicate mean valence and arousal, vertical bars signify 95% confidence intervals for arousal, and horizontal bars represent 95% confidence intervals for valence.

Fig. 1 presents the means and 95% confidence intervals of the affective ratings for the noises and the musical stimuli. Acoustic stimuli varied significantly in both valence and arousal. Excitatory music was interpreted as unpleasant and arousing (valence: −3.08 ± 0.22; arousal: 2.01 ± 0.29). Conversely, the lullaby induced positive emotion with negative arousal (valence: 2.19 ± 0.21; arousal: −1.53 ± 0.28). White noise elicited negative valence (−2.53 ± 0.2) and positive arousal (0.95 ± 0.25). Finally, road traffic noise resulted in negative valence (−1.18 ± 0.23) and arousal near zero (0.5 ± 0.21). The magnitudes of valence were reliably different across the four stimuli (Friedman ANOVA: χ^2^ = 132.46, p < 0.001). This difference was present in pairs of sounds (Wilcoxon: p < 0.05). The levels of sound-induced arousal were significantly different between sound exposures (Friedman ANOVA: χ^2^ = 72.73, p < 0.001). Each sound was different because of the arousal level. This can be appreciated by comparing the effects of the stimuli in Fig. 1.

The analysis results for linear indices of HRV are presented in Table 1.

**Table 1.**
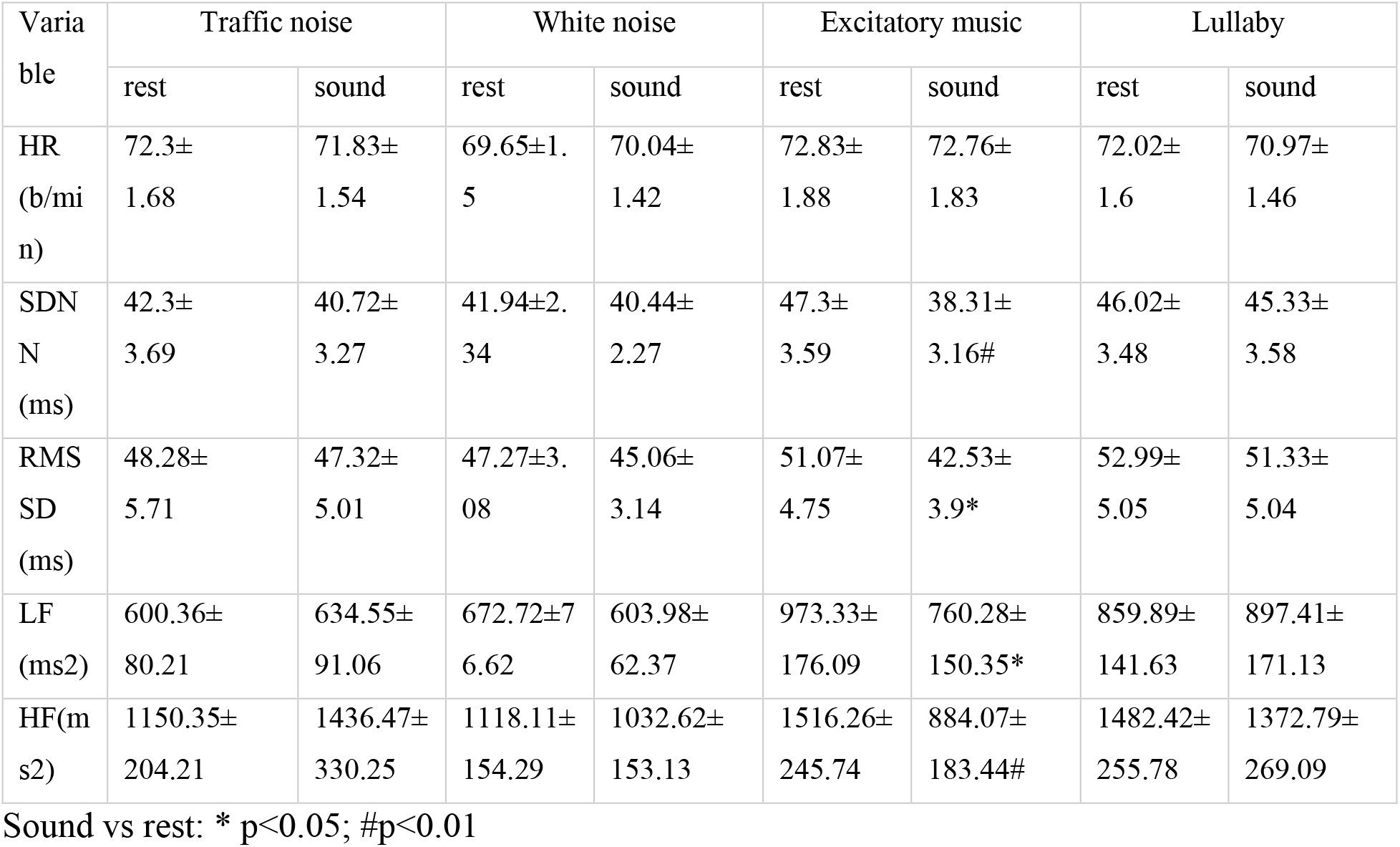
Mean and standard error of time and frequency domain indices during control rest and exposure to auditory stimulation.

We found that sound listening produced small, nonsignificant changes in heart rate (table 1). Excitatory music exposure was associated with a significant reduction in the standard deviation of normal R-R intervals (SDNN). SDNN changes during other sound listening were insignificant. All sounds exposure induced the decrease in RMSSD, but only excitatory music exposure was associated with a significant reduction of vagal activity, as determined by the square root of the mean squared differences of successive R-R intervals.

A decrease in SDNN during excitatory music listening was associated with a significant decline of LF (p<0.05). White noise induces a nonsignificant decrease, traffic noise, and lullaby slightly increased LF (p>0.05). Excitatory music induced a significant decrease of HF (p < 0.01), while the effects of the other sounds were nonsignificant.

The analysis results for nonlinear indices of HRV are presented in Table 2.

**Table 2.** Nonlinear indices of HRV before and during exposure to auditory stimulation.

| Variable | Traffic noise |  | White noise |  | Excitatory music |  | Lullaby |  |
| --- | --- | --- | --- | --- | --- | --- | --- | --- |
|  | rest | sound | rest | sound | rest | sound | rest | sound |
| SD1 (ms) | 33.02±3.55 | 37.3±4.32 | 33.48±2.18 | 31.91±2.22 | 36.16±3.37 | 30.12±2.77* | 37.53±3.58 | 36.35±3.57 |
| SD2 (ms) | 46.5±3.41 | 49.36±3.61 | 48.33±2.74 | 46.81±2.57 | 54.86±4.24 | 45.61±3.61# | 52.34±3.66 | 52.19±3.8 |
|  | rest | sound | rest | sound | rest | sound | rest | sound |
| SD2/SD1 | 1.58±<br>0.07 | 1.51±<br>0.06 | 1.56±<br>0.07 | 1.61±<br>0.07 | 1.64±<br>0.07 | 1.66±<br>0.07 | 1.52±<br>0.07 | 1.55±<br>0.06 |
| SampEn | 1.43±<br>0.04 | 1.41±<br>0.04 | 1.66±<br>0.04 | 1.68±<br>0.04 | 1.4±<br>0.03 | 1.5±<br>0.05 | 1.74±<br>0.03 | 1.77±<br>0.03 |
| D2 | 2.28±<br>0.25 | 2.34±<br>0.24 | 2.74±<br>0.23 | 2.62±<br>0.22 | 2.58±<br>0.25 | 2.14±<br>0.25 | 2.68±<br>0.23 | 2.59±<br>0.22 |
| $\alpha_1$ | 0.86±<br>0.04 | 0.85±<br>0.04 | 0.86±<br>0.03 | 0.89±<br>0.04 | 0.89±<br>0.04 | 0.9±<br>0.04 | 0.83±<br>0.04 | 0.85±<br>0.03 |
| $\alpha_2$ | 0.32±<br>0.01 | 0.29±<br>0.01 | 0.29±<br>0.01 | 0.31±<br>0.01 | 0.31±<br>0.02 | 0.32±<br>0.02 | 0.29±<br>0.01 | 0.3±<br>0.01 |

Sound vs rest: \* $p < 0.05$ ; # $p < 0.01$

During the excitatory music listening, the SD1 and SD2 turned lower than the rest (SD1, p<0.05, SD2, p<0.01). The cloud formed by the RRs becomes smaller, indicating lower variability (Figure 3).

**Figure 3.**
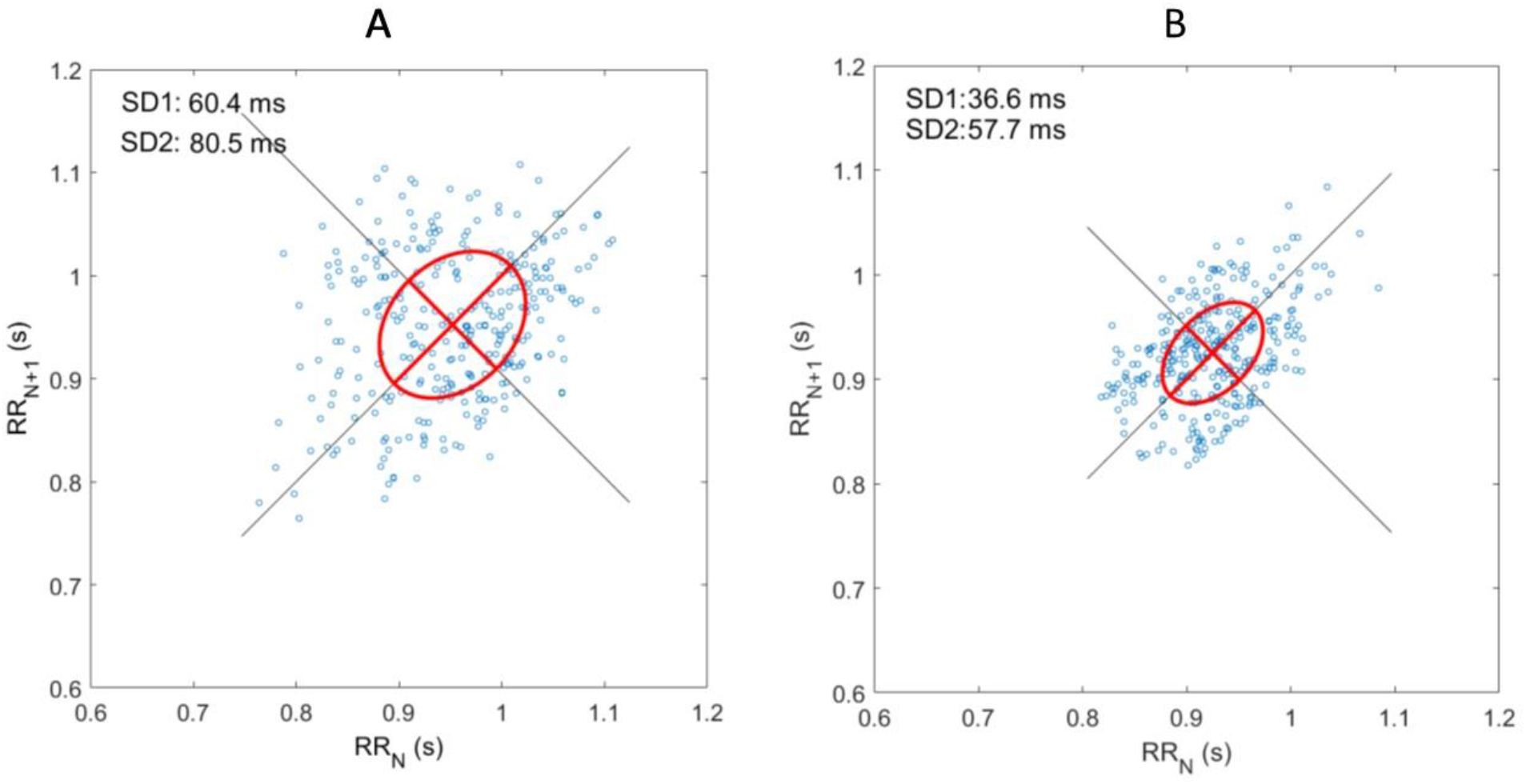
Poincaré plots during rest (A) and excitatory music listening (B).

Analysis of complexity measures (SampEn, and D2) yielded nonsignificant differences between sound listening and rest. Sound listening produced nonsignificant response changes in the shortrange and long-range fractal-scaling exponents of DFA (α1 & α2) (p > 0.05).

Two typical recurrence distances plots during rest and excitatory music listening found in the present cohort and their respective RPs are illustrated in Figure 4.

**Figure 4.**
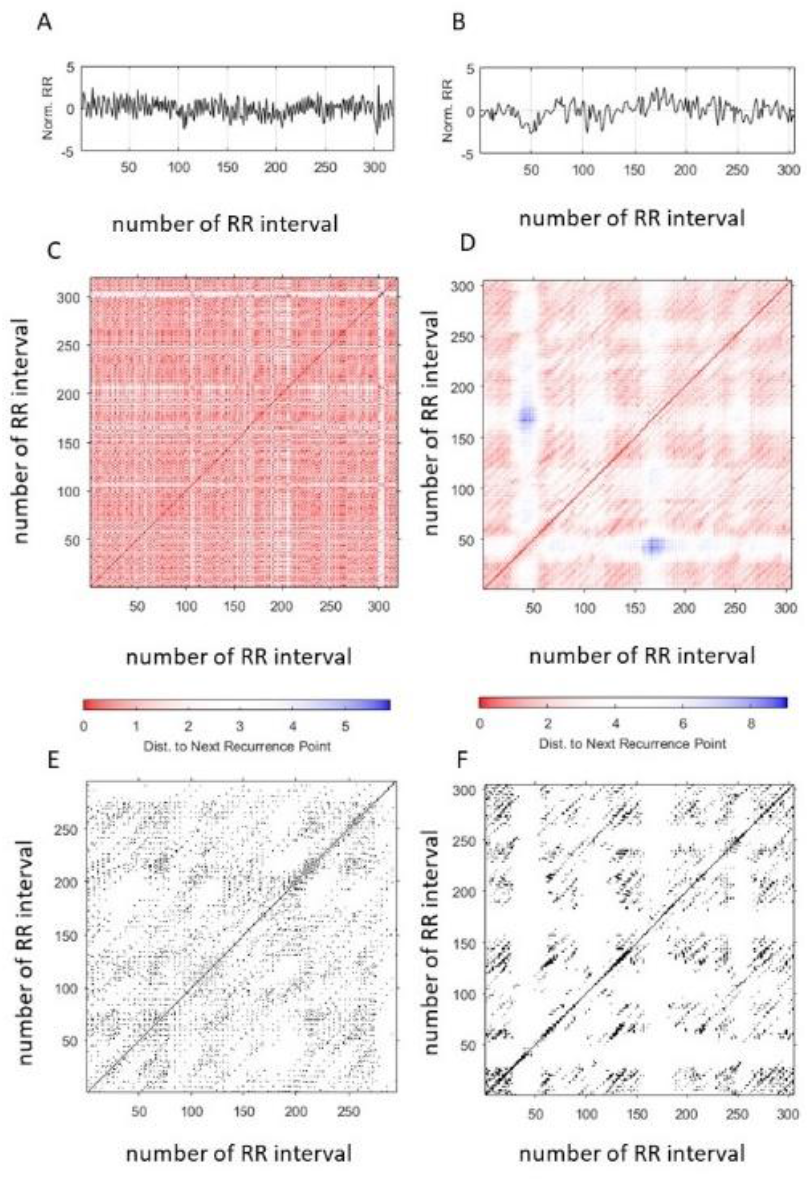
Examples of recurrence plots during rest and excitatory music listening. The top of the figure depicts the RR series for rest (A) and sound exposure conditions. Unthresholded recurrence plots for output with Euclidean Distance for rest (C) and listening (D). The lower graphs demonstrate the resulting RP of RR intervals for rest (E) and exposure conditions (F) after a threshold ε was applied on the distance plot.

The scary music challenge significantly modified RP. Figure 4 shows an example of RR time series that generate the recurrence plots in rest and listening. During the rest, the high-frequency oscillations of heart rate produce regularly distributed diagonal lines parallel to the identity line. These short diagonal lines are clumped in well-formed squares and rectangles in the distance plot (middle panel, C). Comparing the RR time series and recurrence plots, one can find separate white spaces corresponding to rapid changes of the RR amplitude (e.g., the transient but substantial changes in the RR interval time series that occurred around sample number 303).

Excitatory music induces significant changes in the RR tachogram (figure 4) and a decrease in the high-frequency oscillations, as mentioned above (table 1). As shown in figure 4, the recurrence plot has prominent qualitative changes during excitatory music exposure. Figure 4 (F) shows qualitative changes in diagonal lines (they are bigger and more scattered than the ones observed in rest), and more areas have white vertical spaces in correspondence with transient changes in the RR time series.

The means and standard errors of RQA measures for all sound stimuli are presented in Table 3.

**Table 3.** Results of Recurrence Quantification Analysis.

| Variable | Traffic noise |  | White noise |  | Excitatory music |  | Lullaby |  |
| --- | --- | --- | --- | --- | --- | --- | --- | --- |
|  | rest | sound | rest | sound | rest | sound | rest | sound |
| Lmean<br>(beats) | 0.37±0.<br>03 | 0.37±0.<br>03 | 5.78±0.<br>43 | 5.33±0.4 | 30.4±<br>2.69 | 35.1±<br>3.47 | 4.69±<br>0.28 | 4.25±<br>0.21 |
| Lmax<br>(beats) | 63.33±2<br>0.37 | 51.51±2<br>1.43 | 83.6±20<br>.7 | 69.42±18<br>.09 | 4.12±<br>0.28 | 4.49±<br>0.55 | 61.8±<br>18.68 | 50.31±<br>14.08 |
| DET (%) | 0.39±<br>0.02 | 0.39±<br>0.02 | 0.4±0<br>.02 | 0.44±<br>0.02 | 0.38±<br>0.02 | 0.46±<br>0.02# | 0.38±<br>0.01 | 0.38±<br>0.01 |
| LAM | 0.37±<br>0.03 | 0.37±<br>0.03 | 0.4±<br>0.03 | 0.44±0.0<br>3 | 0.41±<br>0.03 | 0.47±<br>0.03 | 0.38±<br>0.02 | 0.35±<br>0.03 |
| V max | 9.02±<br>0.92 | 9.89±<br>1.17 | 9.67±<br>0.79 | 13.53±<br>2.76 | 10.67±<br>0.93 | 12.76±<br>1.7 | 10.94±<br>1.02 | 9.02±<br>0.83 |
| TT | 2.33±<br>0.05 | 2.39±<br>0.06 | 2.41±<br>0.04 | 2.64±<br>0.16 | 2.38±<br>0.04 | 2.53±<br>0.07 | 2.36±<br>0.04 | 2.32±<br>0.05 |
| ShanEn | 1.47±<br>0.12 | 1.32±<br>0.1 | 2.41±<br>0.04 | 2.64±<br>0.16 | 1.18±<br>0.11 | 1.27±<br>0.12 | 1.43±<br>0.11 | 1.28±<br>0.1 |

Sound vs rest: \* $p < 0.05$ ; # $p < 0.01$

The RQA indices related to the diagonal lines (Lmean and Lmax), indicate the absence the significant changes induced by sound exposure. At the same time, we discovered that exposure to excitatory music significantly increases the ratio between recurrence points that form diagonal structures (DET) associated with deterministic behavior. In addition, we found no change in ShanEn, which reflects the dispersion in the probability distribution of the diagonal line lengths.

## Discussion

This study purposed to analyze the effects on HRV of auditory stimulation by different sounds and music styles, using the time-domain, frequency-domain, and nonlinear measures. It is well known that music evokes a wide range of basic and complex emotions. Indeed, such emotional responses play a key role in music perception and autonomic response to affective sounds [46; 47]. In this study, affective sounds evoked negative emotions with different arousal levels and positive emotions with moderate arousal. According to the literature, the autonomic nervous system is an essential part of the emotional response to visual and auditory stimulation, so we hypothesized that sounds used in our experiment might induce significant changes in HRV parameters. Nonsignificant changes in HR contradict conventional expectations, that strong negative emotions evoke an increase in HR [7]. We found considerable variability in individual heart rate changes during sound listening consistent with early published studies. Ivonin et al. [48] found that unpleasant audio stimuli induce HR deceleration, and Orini et al. [49] reported a significant decrease in RR intervals while listening to unpleasant music.

SDNN reflects all the cyclic components of heart rate variability during sound exposure; the primary source of the SDNN is parasympathetically-mediated respiratory sinus arrhythmia (RSA) [50]. Our results are consistent with earlier findings [51] and suggest that music-induced high-arousal negative emotions cause inhibition of SDNN. According to literature, listening to relaxing music lowers sympathetic activity and causes excitement in the parasympathetic branch of ANS [52]. Lullaby music listening induced an insignificant decrease in SDNN, consistent with the effects of live music therapy among hospitalized women [53]. Traffic noise exposure (70 dB(A)) exerts a nonsignificant decrease in SDNN. This result may be explained by the inverse U-shape relationship between equivalent continuous sound pressure level and SDNN [54]. Kraus U et al. [54] found that 5-min daytime exposure of noise ≥ 65dB(A) induces nonsignificant changes in SDNN.

The results show that RMSSD, a measure of parasympathetic control of the heart rate, significantly decreases during excitatory music listening. The insignificant response of RMSSD to arousal music (heavy metal) was reported in several studies [55, 56]. We hypothesized that a prominent reduction in vagal activity in response to Women with Steak Knives might be explained by the negative valence of induced emotions [57].

Low-frequency power reflects both sympathetic and parasympathetic activities [9], and music-induced emotional arousal results in sympathetic nervous system activation [58]. That allows us to assume that associated with high valence and negative arousal decline of LF was due to decreased respiratory sinus arrhythmia (RSA). This assumption was confirmed by a decrease in HF during excitatory music listening. Effects of other sound exposure (white noise, traffic noise, lullaby) on LF were insignificant. The analysis of relevant literature reveals contradictory empirical findings: while most HRV studies of music listening demonstrate nonsignificant changes in LF during listening relaxing and arousal sounds [53, 56, 59], several studies report increased or decreased LF levels during the listening of calm or arousal sounds [51, 60]. RSA was unaffected by white noise, traffic noise, and lullaby listening, evidenced by nonsignificant changes in HF. In previous studies relaxing baroque music-induced significant decrease [56, 51] or nonsignificant changes in HF [61], heavy music did not affect this measure of HRV [59] or significantly reduced it [51]. Our results confirm vagal withdrawal during an arousal sound exposure and challenge the assumption regarding the increase in RSA by relaxing music listening.

In the Poincaré plots, the SD1 and SD2 decreases during excitatory music listening, and SD1/SD2 ratio remains unchanged with a concurrent change of shape. Previous studies have demonstrated that Poincaré parameters, SD1 and SD2, may primarily reflect parasympathetic functioning [62]; therefore, changes in Poincaré plot morphology, associated with negative valence and high arousal, appear to be a manifestation of parasympathetic withdrawal, rather than sympathetic activation. A study performed by da Silva et al. [51] showed that SD2 is negatively influenced by excitatory heavy metal music. Another study showed that auditory stimulation with heavy metal music did not affect the shape of the Poincare plot [55]. Unfortunately, the authors did not report the valence and arousal of emotional experience (evoked by heavy metal music), but it was assumed that excitatory music caused more pronounced negative emotions.

Our results indicate that sound exposure led to nonsignificant changes in SampEn. Indices of SampEn, as the indicator of nonlinear heart rate dynamics, evaluate the inherent complexities of RR interval time series under different emotional states (neutral, fear, sadness, happiness, anger, and disgust), evoked by video [63]. However, data about SampEn changes due to music-associated emotions are scarce or non-existent. It is not completely clear how music listening affects complexity and regularity measured by SampEn. Pérez Lloret et al. [64] noted that “New age” music induces a decrease in SampEn, whereas listening to other relaxing music selected by each subject showed no significant effect of music on the HRV parameter. This research conducted the first comparative study influence of different sounds on SampEn.

This study did not find that sound evokes changes in the complex structure of the attractor, as evidenced by a lack of significant differences in D2 between rest and listening sessions. Vanderlei et al. [65] investigated the effect of heavy metal music and “Kinderszenen” by Robert Schumann on the correlation dimension of HRV and established the absence of statistically significant difference between musical stimulation and control conditions.

DFA is a widely used technique for detecting fractal dynamics in noisy, nonstationary physiological signals and, in the case of HRV analysis, provides information about long-range correlations of RR intervals. There are conflicting data in the literature about the sensitivity of DFA indices to sound exposure, with some data suggesting a significant effect [65, 66]. We find no evidence that sounds induce significant changes in short- and long-term exponent (termed α1 and α2).

We investigated the dynamic structure of cardiac interbeat intervals collected during exposure to silence or sounds in this work. The results of the research indicate that heart rate dynamics were not completely deterministic (DET <1), but a deterministic structure was present in the heart rate variations (DET was statistically >0). Determinism was significantly higher for excitatory music exposure than for silence, suggesting the transition from stochastic to deterministic heart rate behavior. In Chen et al. [68] study, the low-frequency noise-induced significant increase in the length of the longest diagonal line (Lmax) and laminarity (LAM) RQA. RQA features (TT, LMAX, DET, TT) changed significantly during the listening of motivational song, suggesting the shift in the cardiac activity due to the sound exposure [69]. Augmentation of DET generally displays a more frequent return of the heart rate to previous states]. These observed changes in the recurrence measures of heartbeat activity might mirror the increase of constraints on heartbeat variability during excitatory music listening. Increase in DET can be interpreted as a stronger attractor for heartbeat fluctuations, created by negative valence and high arousal emotion.

White noise is an example of an unnatural sound, where similar sound intensities are delivered across a wide frequency range [70]. White noise cannot naturally occur and is unrelated to natural life events. White noise is often reported as aversive stimuli that shift the autonomic balance to sympathetic predominance [71]. Our examination of autonomic effects has found nonsignificant changes in vagal activity, consistent with previously published data [59].

Participants interpreted traffic noise as background sounds and exhibited a weak emotional response to this stimulus. Our results run counter to the stereotype that traffic noise induces a shift in autonomic balance toward sympathetic predominance with vagal withdrawal [17]: noise exposure-induced changes in time domain and frequency domain measures were insignificant. These findings are consistent with previous studies in which no significant effects of traffic noise on linear measures of HRV [66]. The absence of effect may be explained by the low noise exposure level and low emotional response to the sound.

The experiment was designed to induce a wide range of emotions, from activated negative to unactivated positive. The excitatory music with negative valence evokes significant withdrawal of vagal activity and reduction in heart rate complexity, which is quite expected and can be explained by activation of limbic and paralimbic neural circuitry [72]. Although it is believed that listening to relaxing music reduces cortisol levels, decrease heart rate, and mean arterial pressure [53], the results show the absence of significant changes in cardiac autonomic activity, assessed by HRV measures. Our results are consistent with a previous study that already showed nonsignificant changes in HRV parameters during the listening of relaxing music [73].

The single isolated relatively low level of sound in this study is the main limitation that makes it impossible to study the relationship between sound pressure level and effect. On the other hand, we have revealed the apparent effect of emotions evoked by the sound since more loud sounds (100 dB) elicit the autonomic response, mediated by activation of the vestibular system [74]. We did not find a significant difference in heart rate response to sounds with different valence. These results are consistent with the results of the previous study [60]. An exciting feature of our results is the dissociation between the insignificant decrease of HR and significantly reduced respiratory sinus arrhythmia. The dissociation between HRV and HR indicates that HRV is a more sensitive autonomic measure of sound-induced emotions.

### Conclusion

In conclusion, our study indicates that prominent negative emotions, induced by excitatory music, cause the significant vagal withdrawal. Negative emotions in music evoke prominent decrease in short-term and long-term variability of RR intervals in first-return Poincaré plots. This study has shown that measures derived from recurrence plots can provide additional information about the nonlinear complexity of human cardiovascular control systems during affective sound listening.

